# Evolutionary dynamics of male reproductive genes in the *Drosophila virilis* subgroup

**DOI:** 10.1101/111377

**Authors:** Yasir H. Ahmed-Braimah, Robert L. Unckless, Andrew G. Clark

## Abstract

Postcopulatory sexual selection (PCSS) is a potent evolutionary force that can drive rapid changes of reproductive genes within species, and thus has the potential to generate reproductive incompatibilities between species. Male seminal fluid proteins (SFPs) are major players in postmating interactions, and likely the main targets of PCSS in males. The virilis subgroup of *Drosophila* exhibits strong interspecific gametic incompatibilities, and can serve as a model to study the genetic basis of PCSS and gametic isolation. However, reproductive genes in this group have not been characterized. Here we use short-read RNA sequencing of male reproductive organs to examine the evolutionary dynamics of reproductive genes in members of the virilis subgroup: *D. americana, D. lummei, D. novamexicana*, and *D. virilis.* For each of the three male reproductive organs (accessory glands, ejaculatory bulb, and testes), we identify genes that show strong expression bias in a given tissue relative to the remaining tissues. We find that the majority of male reproductive transcripts are testes-biased, accounting for ~15% of all annotated genes. Ejaculatory bulb-biased transcripts largely code for lipid metabolic enzymes, and contain orthologs of the *D. melanogaster* ejaculatory bulb protein, Peb-me, which is involved in mating-plug formation. In addition, we identify 71 candidate SFPs, and show that this set of genes has the highest rate of nonsynonymous codon substitution relative to testes- and ejaculatory bulb-biased genes. Furthermore, these SFPs are underrepresented on the X chromosome and are enriched for proteolytic enzymes, which is consistent with SFPs in other insect species. Surprisingly, we find 35 *D. melanogaster* SFPs with conserved accessory gland expression in the virilis group, suggesting these genes may have conserved reproductive roles in *Drosophila.* Finally, we show that several of the SFPs that have the highest rate of nonsynonymous codon substitutions reside on the centromeric half of chromosome 2, which contributes to paternal gametic incompatibility between species. Our results suggest that SFPs are under strong selection in the virilis group, and likely play a major role in PCSS and/or gametic isolation.

## Introduction

In sexually reproducing organisms, the ability to secure mates is a central component of fitness. Male mating success is largely determined by behavioral traits that are often under sexual selection. Furthermore, in species where females store sperm for extended periods and mate with multiple males, sperm can compete for fertilization and/or females can bias fertilization to certain males [10]. This additional layer of sexual selection—known as postcopulatory sexual selection (PCSS)—can drive rapid evolution of genes involved in postcopulatory interactions. Indeed, reproductive genes evolve rapidly in many animal taxa, often by positive selection [48]. Importantly, rapid evolution of reproductive genes can have direct consequences for speciation by establishing barriers to fertilization between divergent populations [33]. The pattern of rapid evolution of reproductive genes and the potential involvement of PCSS in this divergence is widely recognized. However, the molecular genetic basis of PCSS is still not well understood [56].

Among internally fertilizing organisms, a complex interaction takes place between the female reproductive tract and the male ejaculate, ultimately leading to the union of female and male gametes [58]. Extensive studies in *Drosophila melanogaster* reveal that these interactions are mediated in part by seminal fluid proteins (SFPs), which are secreted from the paired male accessory glands (AGs) and the ejaculatory bulb (EB) [41]. Some of these proteins induce physiological effects in the mated female, such as increased oviposition [22], facilitation of sperm storage [34], reduction of the female’s propensity to remate [50], and reduction of her lifespan [12]. To date about 200 SFPs have been identified in *D. melanogaster,* and several of these have been shown to evolve rapidly between species of the melanogaster group [9,47]. Because of this rapid evolution, only a subset of these genes have orthologs in distantly related species [21]. This pattern suggests that the group of reproductive genes may differ in content across different taxa [25]. Thus, a comprehensive understanding of postcopulatory interactions and their consequences for speciation would benefit from genetic studies on reproductive genes from a wide range of taxa [55].

Closely related species that exhibit gametic incompatibilities in interspecific crosses—often called postmating prezygotic (PMPZ) reproductive isolation—provide a unique opportunity to study the genetic basis of PCSS, because the genes that underlie these gametic incompatibilities likely diverge through PCSS mechanisms within species. Genetic mapping is the traditional approach to identify regions of the genome that cause reproductive isolation between species, and the repertoire of reproductive proteins that reside in these mapped regions represent strong candidates for gametic isolation between species, as the incompatible components in PMPZ necessarily involve the male ejaculate and the female reproductive tract. Male reproductive proteins, particularly SFPs, have been characterized in several taxa, including mosquito [46], honeybee [7], mouse [14], and *D. melanogaster* [17,53]. Many other taxa that are amenable to detailed genetic study and display interspecific PMPZ phenotypes, however, remain largely uncharacterized [55].

Several *Drosophila* species groups provide excellent material for the genetic study of prezygotic reproductive interactions [33]. One such species group is the virilis group (Figure S1A). Members of this group evolve gametic incompatibilities rapidly [3,23,24,44,49]. A subset of virilis group species (the *D. virilis* subgroup: *D. americana, D. novamexicana, D. lummei,* and *D. virilis*) show strong gametic incompatibilities in nearly all heterospecific hybridizations (Figure S1B). In particular, five of the six heterospecific cross combinations between these species produce <2% hatched eggs in at least one direction of the cross [2, 3, 44, 49]. The only species cross in which both directions show appreciable hatch rates is the cross between *D. lummei* and *D. virilis* (∼ 40% fertilization success). The reason for this general reduction in hatch rate was studied in three of these crosses, and appears mostly due to defects in sperm storage [2, 3,44], but other postcopulatory defects are likely. Genetic analyses also reveal that the genetic architecture of the incompatibility between *D. americana* males and *D. virilis* females is somewhat complex: at least three regions on the centromeric half of chromosome 5 (Muller C) and a large inversion on chromosome 2 (Muller E) carry genes responsible for the paternal component of PMPZ [2,49].

Divergent reproductive genes are likely involved in gametic incompatibilities in the virilis group. The processes disrupted in heterospecific inseminations largely resemble those for which SFPs have been implicated in *D. melanogaster* [58]. However, almost nothing is known about reproductive genes in the virilis group. Furthermore, given that *D. virilis* is ∼40 million years divergent from *D. melanogaster,* the virilis group is likely to contain a unique and/or differentiated complement of SFPs.

Here we use short-read RNA sequence data from male reproductive tissues of the *Drosophila virilis* subgroup, in addition to whole-genome sequence data, to characterize the repertoire of reproductive genes in this species group. We obtain RNA-seq data from the accessory glands, ejaculatory bulb and testes (Figure 1). These tissues comprise the main sources of ejaculate components, i.e., sperm and seminal fluid. We also obtain RNA libraries from the gonadectomized male carcass to identify reproductive tissue-specific transcripts. Our objectives are to (1) identify candidate SFPs and tissue-biased reproductive genes, (2) identify functional categories that are enriched among reproductive genes and their potential biochemical roles, (3) examine gene expression and sequence divergence among reproductive genes between species, and (4) identify the set of conserved SFPs between the virilis subgroup and *D. melanogaster.* While we present analyses on all three reproductive tissue types, we focus largely on SFPs, as we think they are the main actors in postmating interactions within the female, and likely the main targets of PCSS in males.

**Figure 1.**
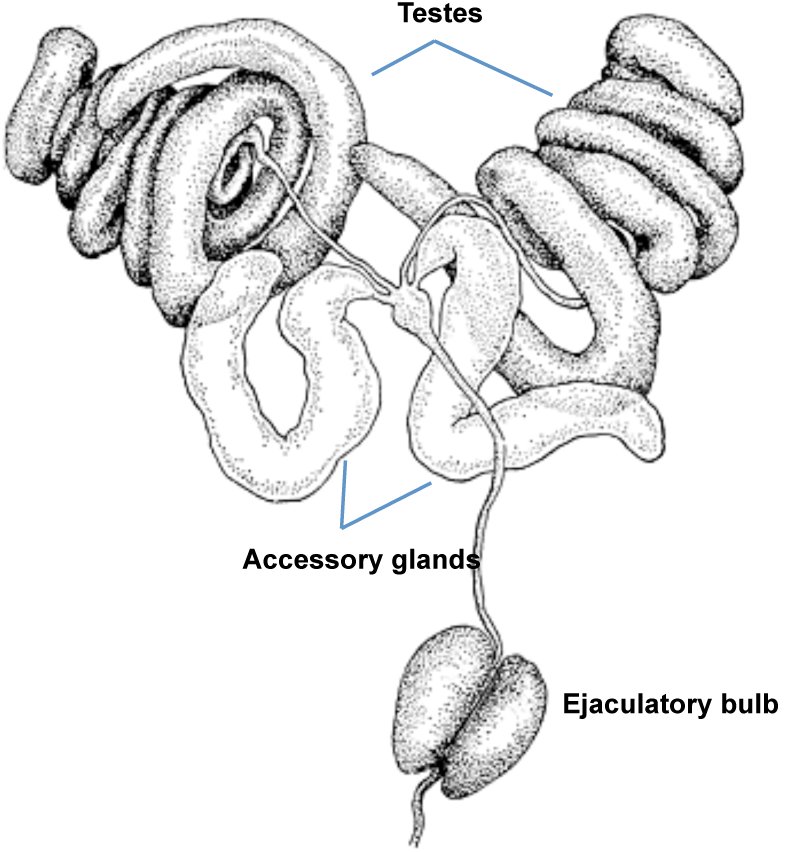
*D. virilis* male reproductive organs [37]. Obtained with permission from FlyBase.

We show that several SFPs are rapidly evolving in this species group, as revealed by an excess of nonsynonymous substitutions relative to other reproductive gene classes and the genome average. Interestingly, the most rapidly evolving SFPs reside within an inverted region that has been implicated in PMPZ isolation between species. We also find that several SFPs show confined expression to one or more species, suggesting that expression divergence is driving differentiation in SFP content. Furthermore, we show that enriched functional categories of SFPs are largely similar to those from other known insects, where proteolytic enzymes dominate. Finally, we identify an appreciable number of conserved SFPs between *D. melanogaster* and the virilis subgroup, suggesting deep conservation of a subset of male ejaculate components.

## Materials and Methods

### Fly strains and hatch rate estimates

Flies were maintained at a constant temperature (22°C) in a 12-hr day/night cycle on standard cornmeal media. A single strain each of D. *virilis* (1051.87), D. *americana* (ML97.5), *D. lummei* (LM.08) and *D. novamexicana* (15010-1031.04) were used throughout this study. Conspecific and heterospecific hatch rate estimates for all possible cross combinations were obtained by calculating the percentage of hatched eggs from daily collections (∼10 collections) of 100 eggs from population cages containing ∼400 males and females. Mean hatch rates represent the average of multi-day collections.

### Tissue dissection, RNA library preparation and sequencing

Individual 12-14 day old virgin males and females of each species were paired and allowed to mate overnight. Males were then anesthetized with CO2 and their reproductive tissues removed. The accessory glands (AG), ejaculatory bulb (EB) and testes were separated and flash-frozen in liquid nitrogen. Each sample consisted of two (EB) or three (AG, carcass, and testes) replicates, and each replicate contained tissue from ∼ 20 flies. RNA was isolated using the Qiagen RNeasy Mini Kit (cat. no. 74104). Paired-end and single-end libraries were prepared using the Illmuina TruSeq RNA Library Prep Kit v2 (www.illumina.com). Sequencing was performed in two stages. In the first, paired-end libraries (one replicate each of AG, carcass, and testes) were sequenced on an Illumina HiSeq2500 at the Cornell Biotechnology Resource Center (Cornell University), and single-end libraries (two replicates each of all four tissues) were sequenced on an Illumina HiSeq2500 at the Genomics Resource Center (University of Rochester).

### Whole-genome short read sequencing and assembly

The four virilis subgroup species were used to generate whole-genome sequence data. A pool of ∼20 males and females from each strain were flash-frozen for DNA library preparation. DNA was isolated using the Qiagen DNeasy Blood & Tissue Kit (cat. no. 69504). Paired-end libraries were prepared using the Illumina TruSeq DNA Library Prep Kit (www.illumina.com). Libraries were sequenced on an Illumina HiSeq2500 at the Genomics Resource Center (University of Rochester).

Paired-end reads from *D. americana, D. lummei* and *D. novamexicana* were mapped to the *D. virilis* reference genome (r1.06) using Bowtie2 v2.2.2 [28]. The reference genome was formatted into chromosome arms (Muller elements) using scaffold placement information [45]. Reads were mapped with the “–local” bowtie2 setting, and assemblies were used to extract whole-genome FASTA sequences for each chromosome/scaffold using Samtools 1.3.1 [30] and seqtk (https://github.com/lh3/seqtk).

### Transcriptome assembly

We aligned the RNA-seq reads to the *D. virilis* reference genome (r1.06) using Tophat v2.1.1 [51] with the follwing settings: -N 20 –read-gap-length 3 –read-edit-dist 20. Aligned reads for each sample were assembled using Cufflinks v2.2.1 [52]. Assembly annotations from each sample were merged to produce the transcriptome annotation file.

We checked whether this annotation contains transcripts not present in the r1.06 annotation, and found none. This file was then used to extract whole-transcript sequences (spliced exons) from the *D. virilis* reference genome for downstream analysis. We also assembled *de novo* transcriptomes for each species using Trinity r20140717 [20]. Processed reads from all tissue types within species were pooled and assembled using default parameters.

### Differential expression analysis

To perform differential expression (DE) analyses, we used the genome-based transcriptome to allow comparison across species. (DE results using the *de novo* transcriptomes were virtually identical, but erroneously assembled transcripts complicate the analysis. Thus, we focus on the gnome-based analysis). Reads from each sample were mapped to the transcriptome using Bowtie2 and abundance estimates were obtained using eXpress [42]. Read counts were normalized using the Trimmed Mean of M-values (TMM) method, and a read-count threshold of 200 counts in any given replicate was established as a filter for low-abundance transcripts.

The DE analysis consisted of calculating the fold-change and DE statistics (*p*-value and FDR) either in comparisons between a given tissue sample against remaining samples within species, or between tissue types across species (edgeR, [43]). A transcript was considered significantly differentially expressed if abundance differed between samples by >4-fold with a Bonferroni corrected *p*-value of <0.001. We defined tissue-biased transcripts as those that are significantly higher in abundance in that particular tissue relative to the remaining tissues.

We also used a tissue specificity index (*S*) to describe tissue-bias between species. In particular, we calculated specificity scores among the same tissue across the four species. The tissue-specificity index was calculated using the following formula:

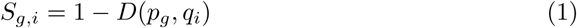
where *D* is the Jensen-Shannon distance, *p_g_* is the expression profile of a given gene *g*, and *q_i_* is a unit for complete specificity in a particular condition *i* [11].

To examine the distribution of tissue-biased transcripts across *D. virilis* chromosomes, we calculated the expected number of tissue-biased transcripts on a given chromosome by multiplying the total number of tissue-biased transcripts in the genome by the proportion of all transcripts on that chromosome. The observed and expected number of tissue-biased transcripts were compared using a χ^2^ test.

### Transcriptome annotation and Gene Ontology (GO) analysis

Genome-based and *de novo* transcriptomes were annotated using several bioinformatic tools. First, predicted open reading frames (ORFs) for the *de novo* transcriptomes were obtained using TransDecoder [20] (ORFs for the genome-based transcriptome were downloaded from FlyBase). Second, mRNA and polypeptide sequences were compared to the Swiss-Prot protein database (www.uniprot.org) using BLASTx and BLASTp, respectively [4]. Third, conserved protein domains, predicted signal peptides, and predicted transmembrane regions were identified using HMMER v.3.1b1 [18], SignalP v 4.1 [38], and tmHMM v2.0 [26], respectively. Finally, Gene Ontology (GO) terms associated with each transcript were extracted from the TrEMBL and SwissProt databases (Trinotate v.2.0). Go term enrichment analyses were performed on sets of candidate tissue-biased genes (GOseq v.1.20.0, [63]).

### Coding sequence divergence

Coding sequence annotations (CDS) of *D. virilis* (r1.06) were used to extract CDS from the whole-genome assemblies of D. *americana*, D. *lummei* and D. *novamexicana* described above (Cufflinks v.2.2.1, *gffread* utility). CDS that contained >20% missing bases (~1% of transcripts) were discarded; >98% of remaining CDS contained less than 5% missing bases. CDS and protein sequences from the four species were aligned using ClustalW v.2.1. Alignments were used to estimate pair-wise *K_a_/K_s_* between species and the mean *K_a_/K_s_* ratio across the virilis phylogeny (ω, PAML v.4.5). In addition, we used PAML’s CODEML program to perform the “branch-site” test on each terminal branch [61]. In particular, the likelihood of a neutral model in which the *K_a_/K_s_* ratio is fixed at 1 is compared to the likelihood of a model in which *K_a_/K_s_* is estimated from the data along each branch of the virilis phylogeny [62]. The test statistic is obtained using the likelihood ratio test (LRT). LRT values were compared to the χ^2^ distribution with 1 degree of freedom. Multiple test correction was carried out by calculating the false discovery rate for each branch class.

### Data and script availability

The raw Illumina reads are available through the Sequence Read Archive under project accession SRP100565. The processed data files and the analysis code are available on GitHub (github.com/YazBraimah/VirilisMaleRNAseq).

## Results

Transcripts that have specialized reproductive roles are likely to have higher abundance levels in reproductive tissues, or even show tissue-specific expression. We compared transcript abundance levels among tissues within species to identify tissue-biased transcripts. We classified these transcripts using a differential abundance level of >4-fold with a *p*-value of <0.001. We used this higher-than-conventional cutoff to be conservative with respect to what is considered tissue-biased in the dataset. With these criteria, we identified 2,493 transcripts that show strong expression bias in the male reproductive tract in all four species (Figure 2). We discuss each tissue-biased gene class below.

**Figure 2.**
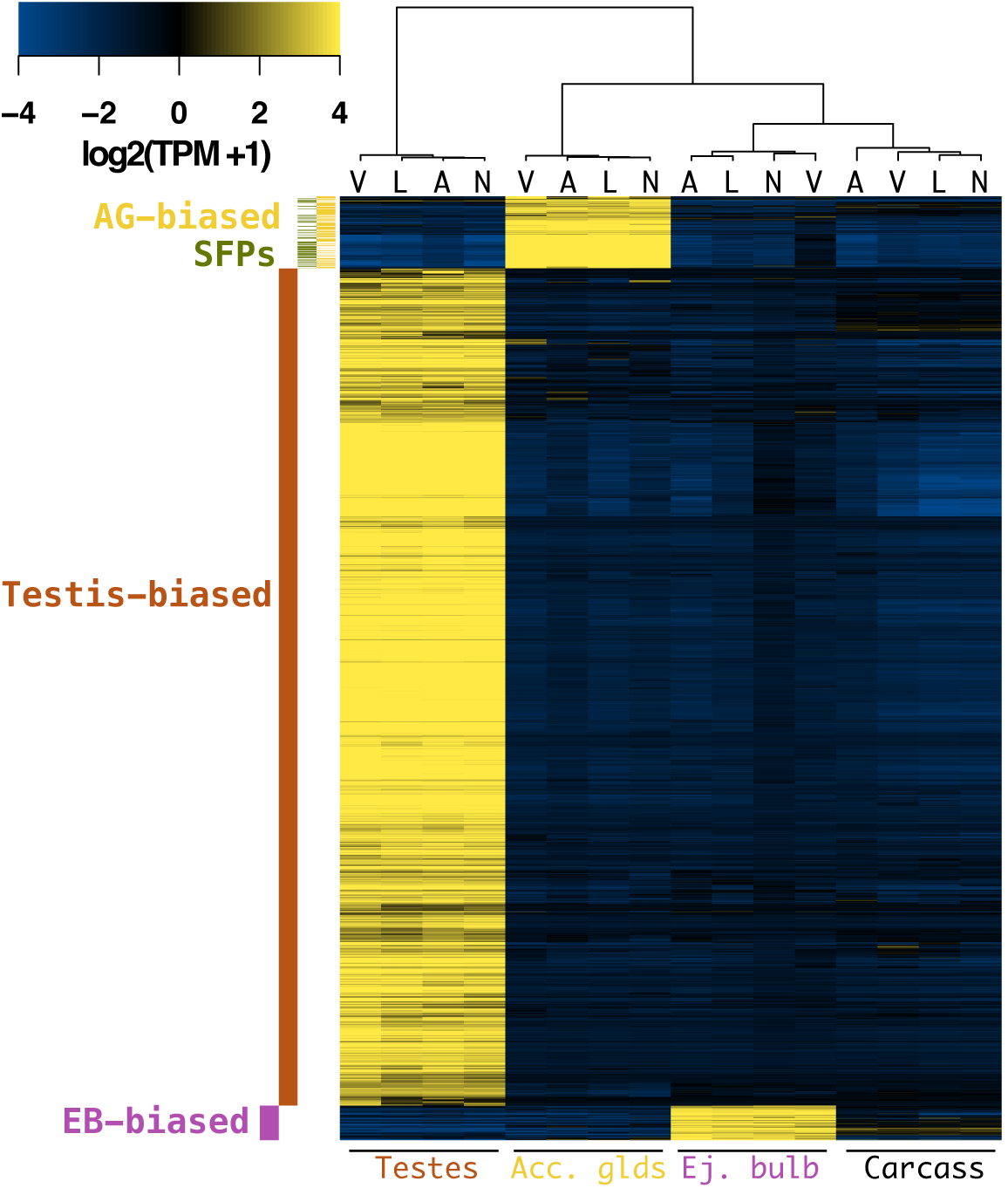
Male reproductive genes of the virilis group: Heatmap of tissue-biased genes that are shared among virilis group members (Heat-scale is median-centered for each gene). The color code on the left indicates the tissue-bias classification (SFPs are the subset of AG-biased genes with a predicted signal peptide). The cladogram on top depicts species clustering (A=D. *americana*, L*=D. lummei*, N=*D*. *novaexicana*, V= *D.virilis*). Species strongly cluster by tissue type, but only clustering of testes-biased genes reflects the true phylogeny.

### AG-biased transcripts and SFPs

The accessory glands are the main source of SFPs, which play an important role in postcopulatory processes [19]. SFPs have also been shown to evolve rapidly in *Drosophila,* and often exhibit lineage-specific gains and losses, most likely through tissue-specific regulatory changes. Here we identified 585 transcripts that show AG-bias in at least one of the four species, but only 191 of these are shared between them (Figure 2, S2). Several transcripts show either species-specific AG-biased expression or are AG-biased in a subset of the four species. Among the AG-biased transcripts that are shared among species, 71 contain predicted signal peptide sequences and are likely to be components of the seminal fluid. Thus, these 71 genes likely play important roles in postmating interactions within the female.

AG-biased genes and SFPs in the virilis group show functional enrichment for several Gene Ontology (GO) categories that have previously been reported for SFPs in other species [7,17,46,58]. Those primarily include extracellular proteolytic enzymes such as serine proteases (Table 1, Figure S1). While SFPs are primarily enriched for extracellular proteolysis proteins and the endoplasmic reticulum, AG-biased transcripts are also enriched for glycosylation enzymes, golgi-associated proteins and carbohydrate-binding enzymes. One example of the latter, *GJ11333,* is an ortholog of the *D. melanogaster* C-type lectin, *Acp29AB,* which has been shown to play a role in sperm storage [59], and shows evidence of positive selection [1].

**Table 1.**
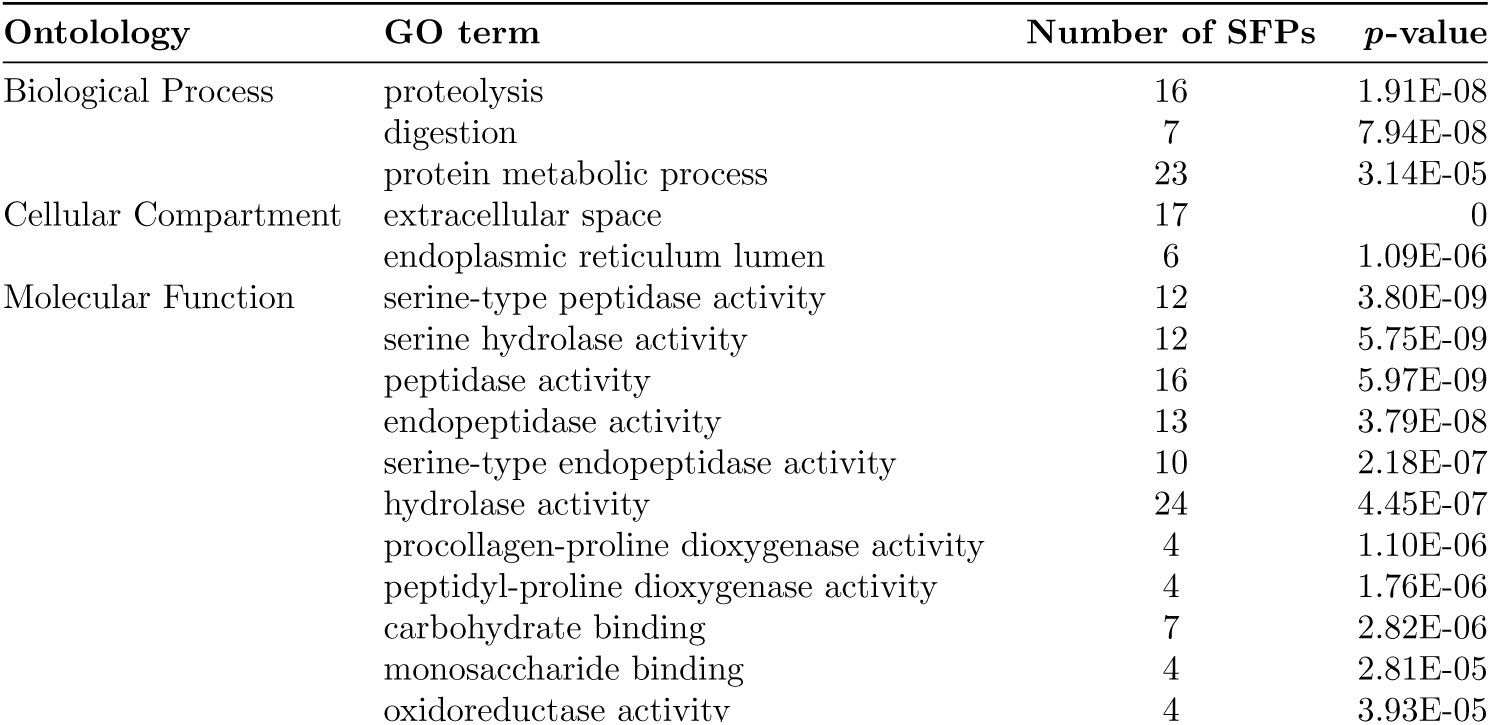
Significantly enriched GO terms among predicted SFPs (*FDR<*0.05).

Male-biased genes are expected to be underrepresented on the X chromosome, particularly if those genes exhibit male-specific reproductive functions [35]. Indeed, we find that AG-biased transcripts and SFPs are not uniformly distributed across the genome (Figure 3). In particular, we calculated the expected number of tissue-biased genes on each chromosome and found that the observed number of AG-biased genes and SFPs is significantly lower on the X chromosome (AG-biased: χ^2^ = 10.1, *p* = 0.002; SFPs: χ^2^ = 8.1, *p* = 0.005). Furthermore, SFPs were significantly enriched on chromosome 2 (χ^2^ = 5.7, *p* = 0.02) and slightly overrepresented on chromosome 4 (χ^2^ = 3.23, *p* = 0.07), while AG-biased genes are significantly enriched on chromsome 4 (χ^2^ = 9.2, *p* = 0.003). The pattern of reduced representation of AG-biased genes on the X chromosome is consistent with findings in *D. melanogaster* [41].

**Figure 3.**
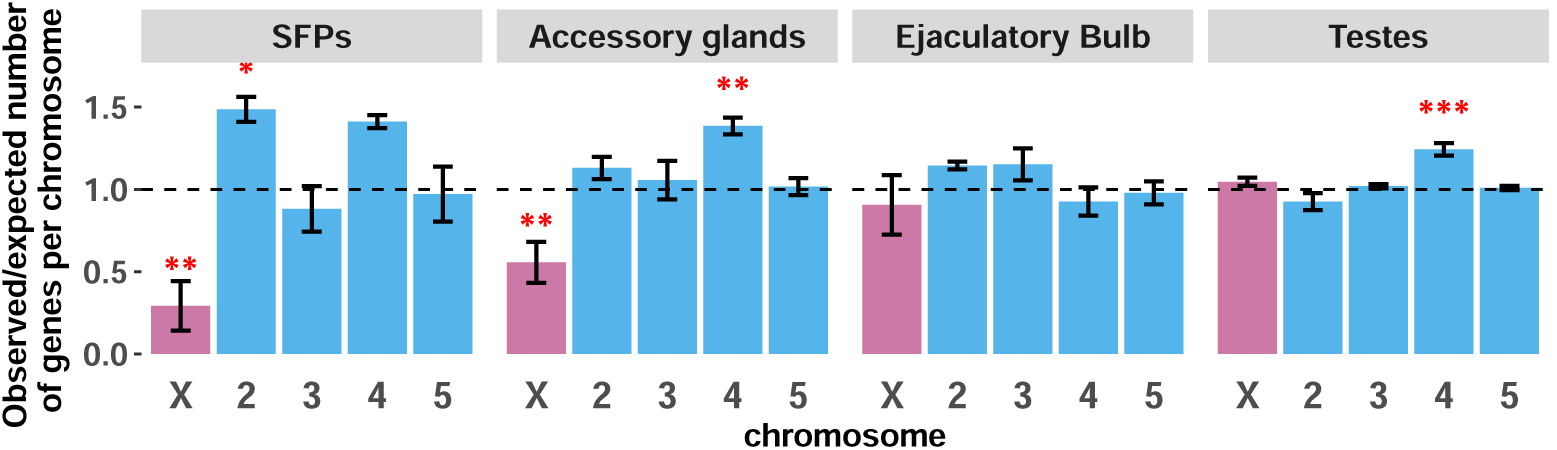
Distribution of tissue-biased transcripts across chromosomes: The ratio of *observed/expected* number of genes on each chromosome averaged between species (dotted line shows random expectation). Significant departures from expectation are depicted by red asterisks (*: *p*<0.05, **: *p*<0.01, ***: *p*<0.001)

In summary, our method identified 71 shared SFPs. These SFPs show functional homology with SFPs in other insect species, and also obey the expected pattern of reduced representation on the X chromosome. Because our approach for identifying SFP candidates is based exclusively on expression abundance and *in silico* prediction of signal peptide sequence, we are likely underestimating the true number of total SFPs.

### EB-biased transcripts

The ejaculatory bulb (EB) is the source of several seminal fluid components in *Drosophila,* most notably the waxy substances found in the copulatory plug [31]. A total of 421 transcripts are classified as EB-biased across the four virilis group species, but only 92 are shared (Figure 2, S2). Of these 92, 20 contain predicted signal sequences, suggesting that these might be components of the ejaculate. Unlike AG-biased genes, EB-biased genes are not significantly underrepresented on the X chromosome (χ^2^ = 0.3; *p* = 0.5, Figure 3).

The EB contributes proteins to the seminal fluid, however the roles of these proteins are poorly understood even in *D. melanogaster.* Some EB-biased transcripts may have similar evolutionary fates as SFPs derived from the accessory glands since they are locked in similar coevolutionary dynamics with females. However EB-biased transcripts are enriched for different functional categories. In particular, EB-biased transcripts are significantly enriched for proteins that are involved in lipid metabolism, fatty acid biosynthesis, steroid metabolism and co-enzyme binding (Table S1), consistent with the likely role of this organ in producing the mating-plug [31]. For instance, a set of six genes are involved in elongation of very long chain fatty acids (*GJ11026, GJ23058, GJ24115, GJ24118, GJ24167, GJ24664*). Another four genes (*GJ19303, GJ21360, GJ22269, GJ26512*) are similar to putative fatty acyl-CoA reductases in *D. melanogaster*.

Finally, we found three EB-biased genes (*GJ20447, GJ21330, GJ22262*) that are orthologous to the *D. melanogaster* ejaculatory bulb proteins, PEB-me and PEB-III. These proteins are integral parts of the mating plug that forms at the vaginal canal after mating [32]. The mating plug is thought to enhance male reproductive success by minimizing sperm loss after copulation and facilitating storage [8]. Thus, some of the genetic components of mating plug formation appear conserved among *Drosophila* species. It is possible, however, that different species contain a different number of copies of Peb orthologs/paralogs.

### Testis-biased transcripts

The largest set of DE genes are those with testis-biased expression. We identified 3179 genes that are testis-biased in at least one of the four species, but 2211 transcripts show testis-biased expression in all four species (Figure 2, S2). Unlike AG-biased genes, testis-biased genes are not underrepresented on the X chromosome (χ^2^ = 0.83; *p* = 0.4, Figure 4), but are significantly overrepresented on chromosome 4 (χ^2^ = 28.2; *p* = 1*x*10^−7^, Figure 4). Genes with testis-biased expression have previously been shown to have a high turnover rate between species [15], and frequently arise on the X chromosome [29].

**Figure 4.**
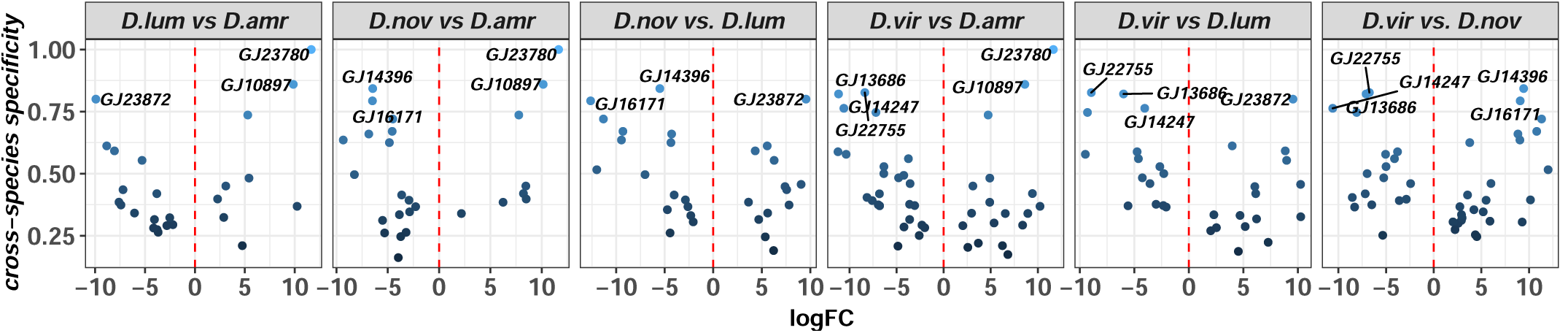
Differential abundance between candidate SFPs across species: Cross-species specificity (*S*) is plotted as a function of fold-change in transcript abundance (log scale) in pairwise comparisons between species (*FDR*<0.001). Transcripts with *S*>0.75 are shown.

Another contrast between testis-biased transcripts and AG-biased genes is the low percentage of testis-biased transcripts that contain signal peptide sequences. In particular, only 6.3% of testis-biased transcripts contain predicted signal peptide sequences, whereas 37.3% and 21.7% of AG-biased and EB-biased genes, respectively, contain predicted signal peptide sequences. This suggests that testes contribute few, if any, secretory proteins to the seminal fluid. This is further reflected in the overrepresented categories of proteins found among testis-biased transcripts (Table S2c). Our analysis shows that testes are significantly enriched for intracellular proteins that are involved in gamete production, microtubule synthesis, mitotic processes, ATP binding and flagellum-mediated motility. These categories highlight the array of genes involved in spermatogenesis and the development of the long sperm tail in these species [39]. In addition, there are many genes (>100) that belong to individual functional categories (e.g., microtubule-based processes, mitotic division, motor activity), indicating high investment in gamete production and sperm energetics by these species.

Production of large numbers of motile sperm is an important component of fitness in sperm competition among insects [36]. The large number of testis-biased genes and their functional enrichment for sperm production and motility terms may reflect this [13,54]. Finally, the high rate of gene turnover among reproductive transcripts may represent a process by which novel genes can be recruited and, ultimately, new beneficial functions may arise. We explore these possibilities below.

### Lineage-specific expression patterns

Lineage-specific genes can arise by changes in gene regulation through tissue-specific cooption of a promoter, or may arise *de novo,* such that a new gene is derived from previously noncoding DNA [64]. Both mechanisms can generate genetic and thus evolutionary novelty. To identify possible lineage-specific transcripts, we used two approaches. In the first approach, we (1) examined differential expression between genes that we classified as tissue-biased in any of the four species, and (2) we calculated the specificity score for each tissue type across all species’ samples. (A high specificity score in this case means that both the tissue and the species show exclusive expression of that transcript). This approach aims to identify transcripts that are divergent at the regulatory level and show tissue-biased expression in only one tissue type. In the second approach, we leverage the individual species’ *de novo* transcriptomes to identify transcripts that are present in the one of the species, but absent in the others.

Using the first approach, we identified a range of lineage-specific expression patterns for reproductive genes, and several cases where genes are expressed exclusively in one species (Figures 4, Figure S3). A significant pairwise difference in abundance (>4-fold, FDR <0.001) among tissue-biased genes and a conservative cutoff for the cross-species specificity index (0.75) can point to cases where genes are exclusively expressed in one species. Indeed, we identify eight SFP candidates that show exclusive expression in one species (Figure 4). Two genes (*GJ10897, GJ23780*) are exclusively expressed in *D. americana,* one gene in D. *lummei* (*GJ23872*), two in D. *novamexicana* (*GJ16171, GJ14396*), and three in *D. virilis* (*GJ22755, GJ14247, GJ13686*). Similar lineage-specific patterns of expression are also observed in the other reproductive tissue-biased genes (Figure S3). Recruiting new genes may be important in modifying a species’ reproductive capabilities, and thus these results provide opportunities for investigating the functional significance of expression divergence, particularly as it relates to reproduction.

In the second approach we sought to determine whether any of the species contain *de novo* transcripts, which we define as being exclusively expressed in one species, and have no ortholog(s) in the other species. We accomplished this by querying tissue-biased transcripts—that we validate as derived from a given species’ genome—against the transcriptomes of the sister species. Under this especially restrictive condition, we find that unique transcripts are rare except in *D. americana* testes and the *D. lummei* ejaculatory bulb, where 35 and 30 transcripts, respectively, have no homology to transcripts of the sister species (Figure S4). The dynamics of losses and gains of reproductive genes among lineages appear complex, and beyond the scope of this current study.

In summary, lineage-specific transcripts that arise by evolutionary changes in gene regulation are common within male reproductive tissue, and while our results indicate that some tissues may experience higher gene turnover rates within species, additional work is required to fully assess the evolutionary dynamic of gene gain and loss.

### Comparison of *D. melanogaster* SFPs to *D. virilis*

The rapid evolution of SFPs in ***Drosophila*** suggests that distantly related species may contain different repertoires of these genes. *D. melanogaster* is ∼40 million years divergent from *D. virilis* and is undoubtedly the species with the best characterized complement of SFPs. We first examined expression conservation between known *D. melanogaster* SFPs and their orthologs in the virilis group; here we relied on orthology assignments described in FlyBase. For the *D. melanogaster* expression data, we used publicly available RNA-seq data from three *D. melanogaster* tissues (AG, testes, head; www.modencode.org).

Of 211 known SFPs in *D. melanogaster* [17], 93 have clear orthologs in the virilis group (Figure 5). The percent identity among orthologous proteins ranges from 20% to 90%, and several *D. melanogaster* SFPs show homology with multiple *D. virilis* transcripts. Of the 93 SFP orthologs in the virilis group, 44 are classified as AG-biased in at least one of the four species, and 35 of those contain predicted signal peptide sequence. The remaining orthologs show various expression profiles, with some transcripts having increased abundance in testes (*n* = 14) and ejaculatory bulb (*n* = 6), and several others (*n* = 40) do not show reproductive tissue-bias. AG-biased transcripts and SFPs in the virilis group that have orthologs in *D. melanogaster* show largely congruent expression profiles with their orthologous SFPs in D. *melanogaster,* suggesting these genes might have conserved reproductive function in males. Unfortunately, the best studied SFPs in *D. melanogaster* are not among these orthologs (e.g. Sex Peptide, ovulin, Acp36DE). Three of the orthologs, however, have been implicated in postmating processes in *D. melanogaster*: (1) *seminase* (*GJ12578* in *D. virilis*), which is a protein that acts in the Sex Peptide network, regulates a proteolytic cascade that affects several postmating processes [27], while (2) *antares* and (3) *aquarius* also act in the SP network to facilitate binding of SP to sperm and transfer of other network proteins [16].

**Figure 5.**
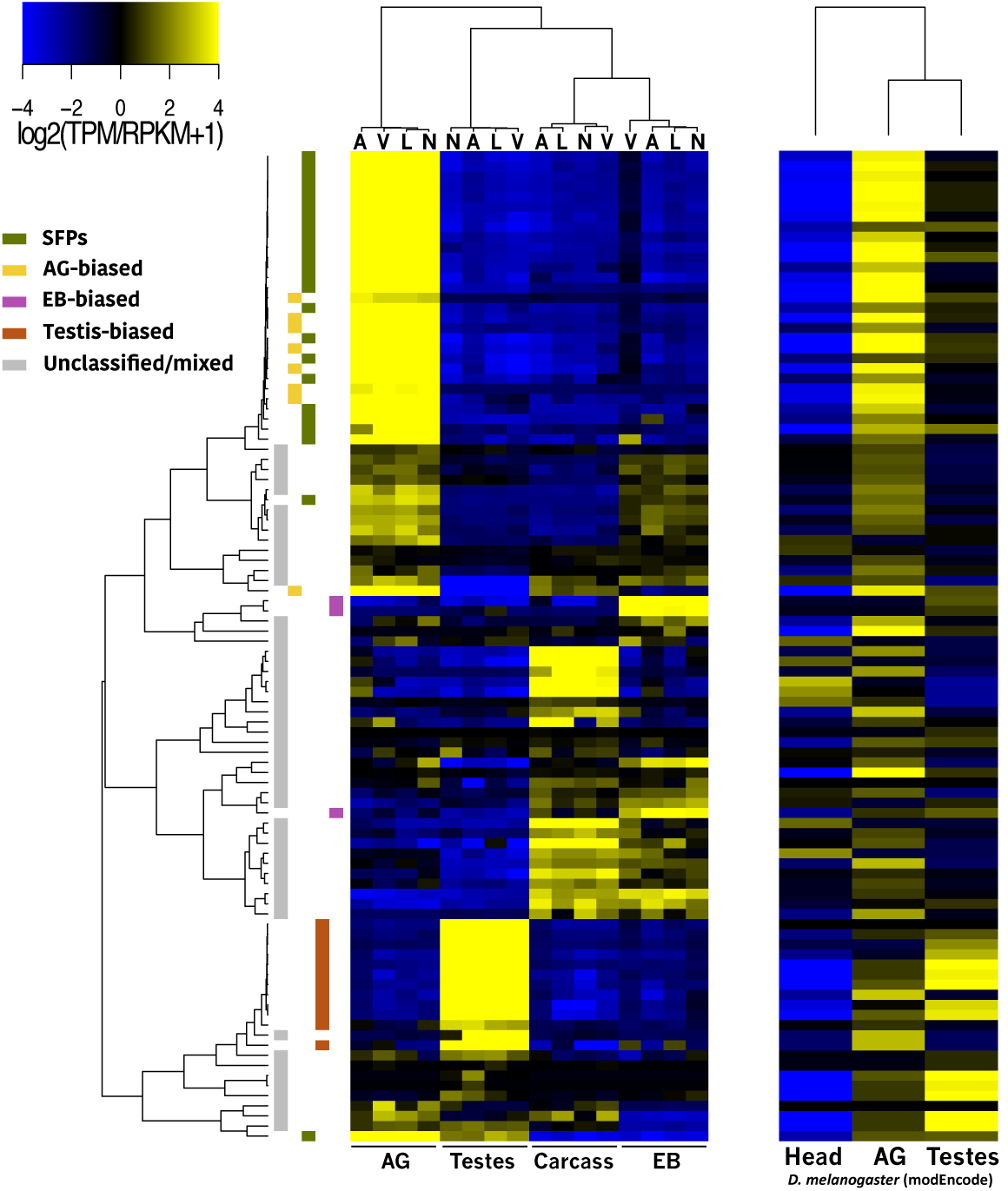
*D. melanogaster* SFPs and their orthologs in *D. virilis*: Expression heatmap of *D. melanogaster* SFPs (right) and their orthologs in *D. virilis* (left). The heat-scale shows expression values in median-centered *log*_2_ TPM (virilis data) or RPKM (*D. melanogaster* data). The left cladogram shows complete-linkage clustering relationships of virilis group orthologs, and the color key on the left indicates shared tissue-bias status among virilis group species. The cladogram on top of the virilis group heatmap depicts species clustering (abbreviations as in Figure 2), and the cladogram ob top the *D. melanogaster* data depicts sample clustering.

We investigated whether the functional classes of *D. melanogaster* SFPs differ from those of AG-biased and SFP transcripts in the virilis group. Most SFPs in *D. melanogaster* are of unknown function [17]. Among *D. melanogaster* SFPs, 29 are known proteases and protease inhibitors. A similar number of proteolytic enzymes are found among *D. virilis* AG-biased transcripts and SFPs (Table 1, S1). Furthermore, several SFPs are classified as lipid metabolism proteins in *D. melanogaster.* These proteins may be products of the EB, as is the case for EB-biased transcripts in the virilis group that are enriched for lipid metabolic processes. It is worth noting that because the identification of SFPs in *D. melanogaster* was performed through proteomic analysis of transferred seminal fluid [17], the expression profiles of SFPs do not always show AG-biased expression in that species (Figure 5, right). Thus, our approach likely misses many proteins that are found in the seminal fluid but do not show AG-biased expression.

These results show that some SFPs are highly conserved between distantly related *Drosophila* species, while others diverge significantly both at the sequence level and at the level of gene regulation. Furthermore, functional classes of SFPs are largely congruent between *D. melanogaster* and *D. virilis*. A more precise comparison would require accurate identification of the transferred complement of SFPs in the virilis group.

### Molecular evolution of male reproductive genes

Reproductive genes can be targets of selection, partly because they play a role in sperm competition or because male and female reproductive genes coevolve as a consequence of cryptic female choice and/or conflicting reproductive interests. Regardless of the mechanism of PCSS, molecular evolutionary analysis of reproductive genes can reveal the impact of such processes on rates of codon substitution and divergence between species. Here we use the ratio of nonsynonymous to synonymous (*K_a_/K_s_*) codon substitutions to (1) examine the difference in average *K_a_/K_s_* across the virilis subgroup phylogeny (ω) for each tissue-biased gene category, (2) examine pairwise *K_a_/K_s_* among SFPs, and (3) test for evidence of adaptive codon substitutions among sites within coding sequences and along branches of the phylogeny using PAML’s branch-site test [62].

First we calculated the mean *K_a_/K_s_* value (ω) across the phylogeny for each tissue-biased category of genes and tested for significant differences from the genome average (Figure 6). All tissue-biased categories deviate significantly from the genome average (ω = 0.15), with EB-biased transcripts showing lower ω (ω = 0.11, W = 1.3*x*10^6^, *p* = 0.02; Wilcoxon rank-sum). AG-biased (ω = 0.3) and testes-biased genes (ω = 0.23), on the other hand, show significantly higher mean ω than the genome average (*W* = 2.9*x*10^6^, *p*<<0.001, and *W* = 3.4*x*10^7^, *p*<<0.001, respectively; Wilcoxon rank-sum). Among AG-biased genes, those we classify as SFP candidates have the highest mean ω (ω = 0.39, *W* = 1.3*x*10^6^, *p*<<0.001; Wilcoxon rank-sum). Thus, our results show that AG-derived proteins—the central players in postmating interactions—experience higher selective pressures than other reproductive gene classes.

**Figure 6.**
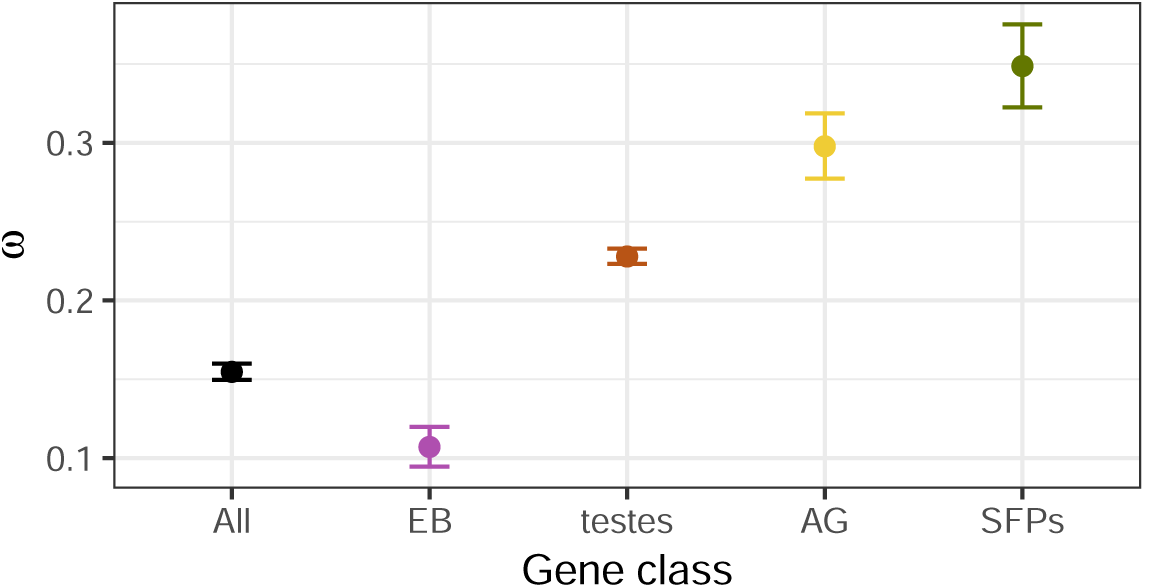
Mean *K_a_/K_s_*(ω) for tissue-biased genes: ω values across the virilis subgroup phylogeny were averaged for each set of shared, tissue-biased transcripts category. The “all” category represents mean ω for all transcripts in the genome. Error bars represent standard error.

Nearly all crosses between virilis group members result in strong gametic incompatibilities (PMPZ), suggesting that PCSS processes within species drove significant differentiation of genes involved in postmating interactions. A striking pattern of PMPZ in this group is the strong incompatiblity between *D. americana* males and females from the sister species. Quantitative trait loci (QTL) that contribute to this paternal incompatibility between D. *americana* and D. *virilis* females have been mapped in two studies [2, 49], and include a large inversion on chromosome 2 and several adjacent QTL on the centromeric half of chromosome 5. We found that the four SFPs with the highest ω values reside within the chromosome 2 inversion (Figure 7). While other tissue-biased categories do not show this pattern, some of these tissue-biased transcripts with elevated ω coincide with PMPZ QTL (Figure S5).

**Figure 7.**
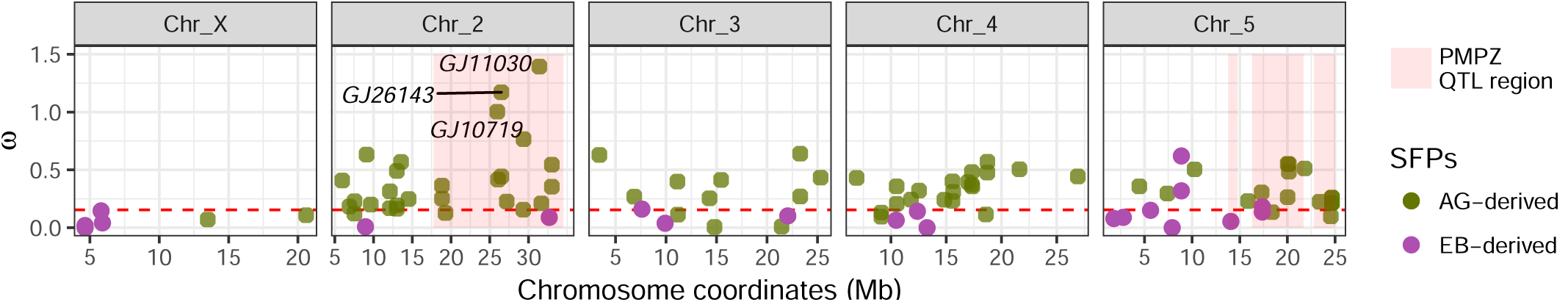
ω values of SFPs along the major chromosomes: ω values for SFPs (AG-derived: green, EB-derived: purple) are shown with respect to genomic location. The shaded pink regions highlight the paternal PMPZ QTLs identified previously (see text). The three SFPs with ω>1 are indicated. The dashed red line indicates the genome average ω.

In the second analysis we examined pairwise *K_a_/K_s_* among SFPs, as some lineage-specific patterns might be missed with ω. Here we find a striking correspondence between SFPs with elevated *K_a_/K_s_* and PMPZ QTL on chromosome 2 (Figure S6). Similar to the ω pattern described above, several SFPs with *K_a_/K_s_>*1 occur on the chromosome 2 inversion. Comparisons involving *D. americana* in particular show 3-4 genes with this pattern, while those comparisons excluding *D. americana* show ≤2 genes. None of the SFPs on the chromosome 5 QTLs have *K_a_/K_s_*>1, but a cluster of 3 elevated genes (*K_a_/K_s_*>0.5) coincide with one of the chromosome 5 QTL. These results suggest that SFPs on chromosome 2 substantially contribute to overall SFP divergence among these species, and particularly along the *D. americana* lineage.

Finally, to test for signatures of positive selection among tissue-biased genes, we performed the branch-site test implemented in PAML [61,62]. This test is conservative, and has higher power to detect positive selection with higher divergence times than the virilis group splits (<5 mya) and a larger number of species in the alignment [5,60]. To perform the test, we calculated the log likelihood of a model where ω=1 (neutral) and compared that to a model in which ω is estimated from the sequence alignment (positive selection). This comparison can be performed using the likelihood ratio test (LRT), where the test statistic under the null hypothesis follows a χ^2^ distribution with 1 degree of freedom. Given that this test is underpowered for our analysis, a signature of positive selection here likely reflects strong adaptive change.

We performed the PAML test on all coding sequences in the genome and calculated the false-discovery rate (FDR) along each branch. We found 92 genes out of 15,078 that show a signature of adaptive evolution (FDR <0.05) along at least one branch of the tree. Three of those are AG-biased, and occur along the *D. lummei* (*GJ13553*), *D. novamexicana* (*GJ17607*), and*D*. *virilis* (*GJ14075*) lineages (Figure S7). A single EB-biased gene (*GJ19792*) contains two rapidly evolving isoforms along the *D. lummei* lineage, and 19 testes-biased genes show a significant signature of positive selection. On the other hand, none of the SFP candidates is significant after multiple test correction, suggesting that selection on these genes is either weak—-few nonsynonymous changes coupled with even fewer synonymous ones (SFPs are generally short)—or undetectable by the LRT.

## Discussion

We identified male reproductive genes in the virilis subgroup using RNA-seq and differential transcript abundance. Using this approach, we provided an overall description of the evolutionary dynamics of these genes in terms of expression divergence, functional classification, distribution across the genome, and sequence evolution. Because these genes play important roles in gametic interactions, understanding their evolutionary dynamics is critical to gain insights into the genetic basis of PCSS and PMPZ.

Testes-biased genes dominate the repertoire of reproductive genes, with >2,000 genes having strong expression bias in that tissue. These genes are highly enriched for GO terms that are linked to gamete development and to basic cell biological processes, often with several hundred genes belonging to individual GO terms. This observation underscores the significant resources that virilis species–and other *Drosophila* species—invest in sperm development. The large number and size of sperm produced by males of these species likely plays an important role in postcopulatory competition between sperm of rival males because, in *Drosophila* and many insects, multiple ejaculates compete for fertilization within the reproductive tract of females [40]. Additional work is needed to uncover the functional importance of the large number of testes-biased genes in affecting PCSS processes.

Ejaculatory bulb proteins are also key players in postmating interactions in *Drosophila,* and some of these proteins play a key role in mating plug formation [32]. The mating plug is a nearly ubiquitous component of male ejaculates in many animal taxa, and is thought to facilitate sperm movement, prevent sperm loss, and prevent subsequent matings [8]. We have identified 92 genes that show strong expression bias in the ejaculatory bulb, and the enriched GO categories among them suggests they are involved in lipid biosynthesis—consistent with their presumed functional roles. In *D. melanogaster,* three proteins (Peb, Peb-II, and Peb-III) play a part in mating plug formation. We identified three genes among virilis species that show homology to two of the *D. melanogaster* Peb genes. This suggests that some aspects of mating plug formation are conserved in *Drosophila.*

The accessory glands are the main source of seminal fluid proteins in *Drosophila,* and many studies in insects have implicated their importance in postmating interactions [6]. In *D. melanogaster,* several of these proteins induce a variety of postmating effects in females [57]. Several SFPs are also known to evolve rapidly among closely related species [47], which suggests that these genes may be the main targets of postcopulatory sexual selection. Because of their rapid evolution, distantly related species may differ in SFP content and/or sequence [25]. It is thus important to independently identify SFPs in species that are distantly related to *D. melanogaster* to gain insights into various genetic mechanisms of PCSS.

Similar to other *Drosophila* species that produce many proteases in the accessory glands [17,25], we found that proteins with proteolytic function are enriched among AG-biased genes and SFPs in the virilis group, suggesting that proteases are a conserved functional class among male seminal proteins in the *Drosophila* genus. We also found that nearly half of known SFPs in *D. melanogaster* have clear orthologs in the virilis group, and nearly a quarter of those show expression conservation. This latter set of highly conserved SFPs across the *Drosophila* genus likely play key roles in postmating interactions. Unfortunately, the majority of these proteins have not yet been characterized in *D. melanogaster.* Three of them, however, were recently shown to act in the Sex Peptide network in *D. melanogaster* by facilitating transfer of other SFPs and binding of SP to sperm [16]. The majority of *D. melanogaster* SFP orthologs in *D. virilis* do not show AG-biased expression among virilis species. These orthologs may have diverged at the expression level and/or have been recruited to other functions. Nonetheless, our general finding of significant similarity between this putatively rapidly evolving class of genes is surprising.

SFPs in the virilis group show the highest rate of nonsynonymous amino acid substitution among the reproductive gene classes. Because SFPs have not been studied in the virilis group, identifying rapidly evolving AG-biased transcripts may reveal interesting candidate genes for further study. The prevalence of gametic incompatibilities as an isolating barrier among members of this group further highlights the utility of investigating rapidly evolving SFPs and their role in PCSS and PMPZ. Indeed, rapidly evolving SFPs in the virilis group are associated with known paternal PMPZ regions. The four SFPs with the highest codon substitution rates reside within a major PMPZ QTL, which is the site of a fixed inversion between *D. americana* and *D. virilis*.

The approach that we have used to identify reproductive genes has several strengths. First, RNA-seq provides information on both RNA abundance and sequence, information that is needed to identify transcripts with tissue-biased expression and to measure rates of nucleotide substitution. Second, by sequencing transcripts from each of the three main reproductive organs from the four species we are able to examine rates of loss/gain of reproductive proteins between species, a phenomenon that can play an important role in reproductive divergence between species. Finally, RNA-seq allows considerable improvement in gene annotations and can identify many new transcripts and splice variants.

Identifying reproductive genes via our approach does have disadvantages. For example, a gene might well play an important role in reproductive processes but not show expression that is specific to reproductive tissues. This limitation can obviously compromise our ability to identify biologically important transcripts. We also used stringent cutoffs in our classification of differentially expressed transcripts to avoid cases of uncertainty due to large variance among replicates, especially with transcripts that have low abundance. Finally, because our inference of secreted ejaculate proteins relies on the presence of signal peptides that is predictable *in silico,* we may miss proteins that are present in the male ejaculate but don’t contain an obvious signal peptide. Despite these caveats, our approach succeeded in identifying many candidate reproductive transcripts for each tissue type, allowing preliminary analysis of such genes in this group.

In summary we have reached several broad conclusions that establish parallels between the virilis and melanogaster species groups in SFP evolution. First, SFPs show an elevated rate of nonsynonymous codon substitution, and that the most rapidly evolving SFPs coincide with known paternal PMPZ QTL. Second, candidate reproductive genes are evolutionarily dynamic such that species may differ in reproductive transcript content, often by regulatory changes. Third, AG-biased transcripts are underrepresented on the X chromosome relative to autosomes. Finally, we identify several orthologs of *D. melanogaster* SFPs, with a subset showing conserved expression patterns suggesting likely functional conservation. Our data and findings provide a powerful platform for further studies of reproductive gene evolution in the virilis group.

## Acknowledgements

We thank Amanda Manfredo for preparing the sequencing libraries, and David Lambert and Daven Presgraves for comments on earlier versions of the manuscript. We especially thank H. Allen Orr for guidance and support throughout this work. This work was supported by National Institutes of Health grants R01-GM051932 to H. Allen Orr and R01-HD059060 to A.G.C.

## Supporting Information

**Figure S1.**
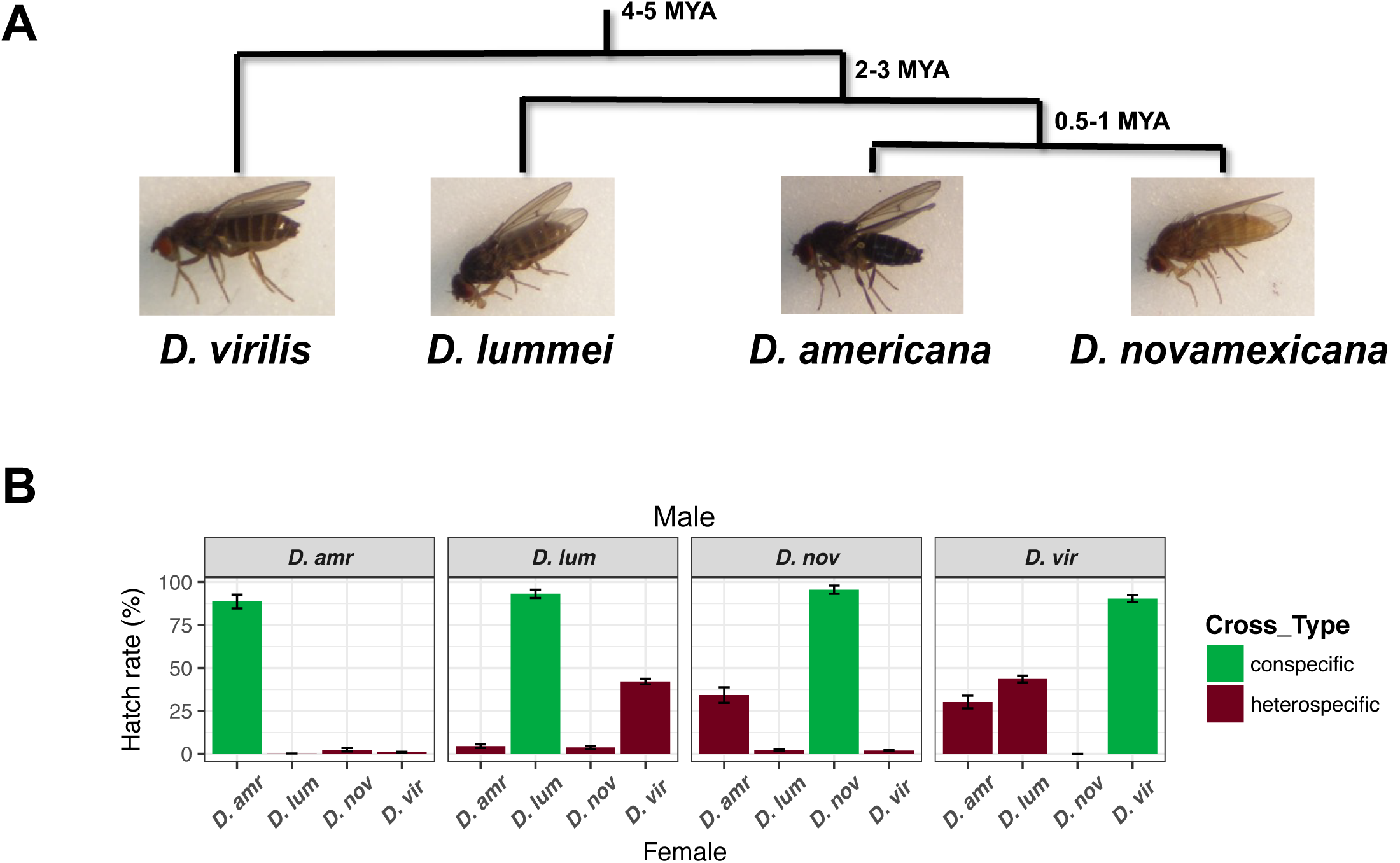
The virilis subgroup as a model for PMPZ and PCSS: (A) Phylogeny of virilis subgroup members. Approximate divergence times are indicated. (B) Mean hatch rate of conspecific (green) and heterospecific (brown) crosses between members of the virilis group. Male genotypes are shown on the gray strip, and female genotypes on the *x*-axis. Error bars represent standard error.

**Figure S2.**
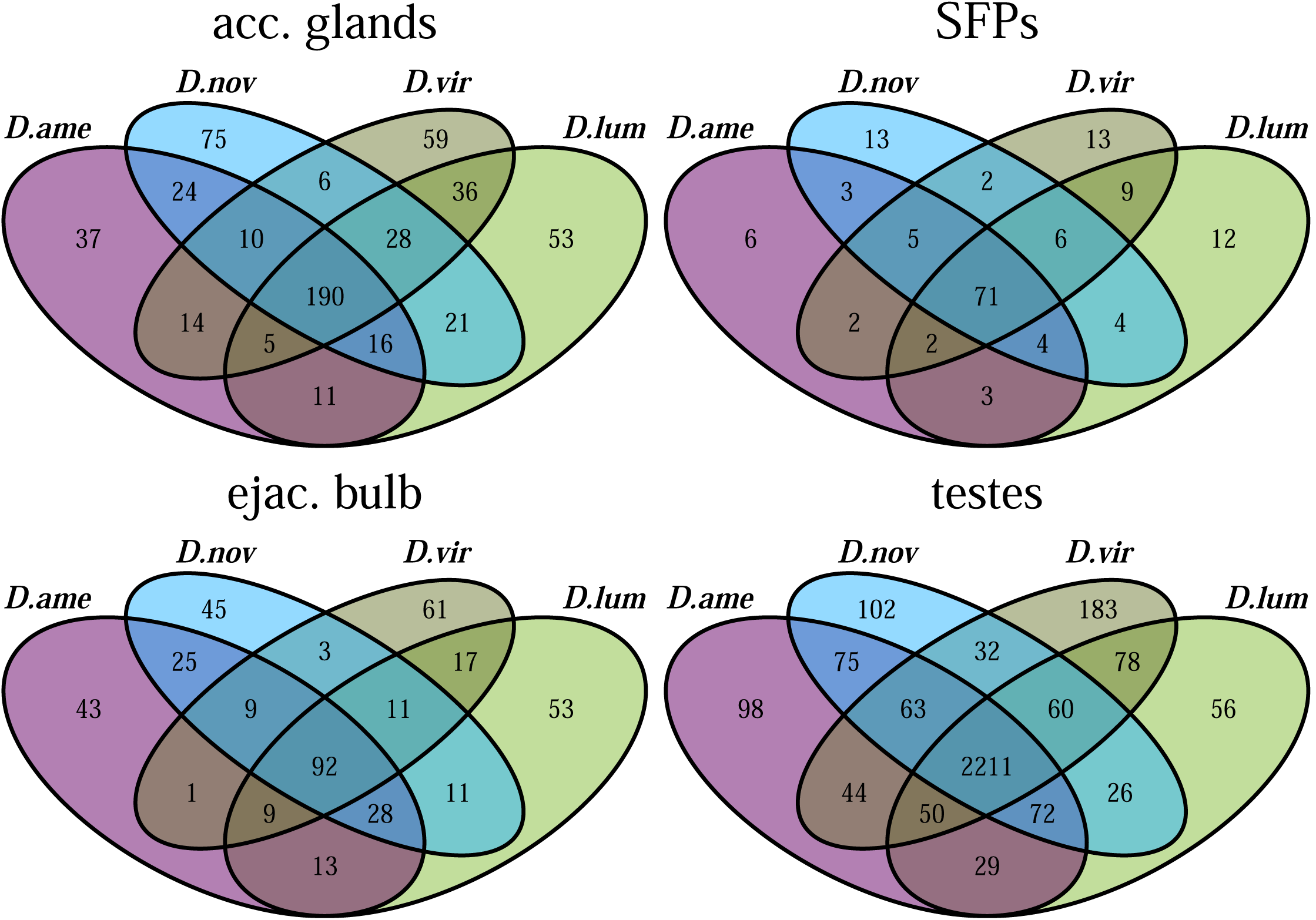
Tissue-biased transcript numbers: Venn diagram of tissue-biased transcripts (FDR <0.001) grouped by species.

**Figure S3.**
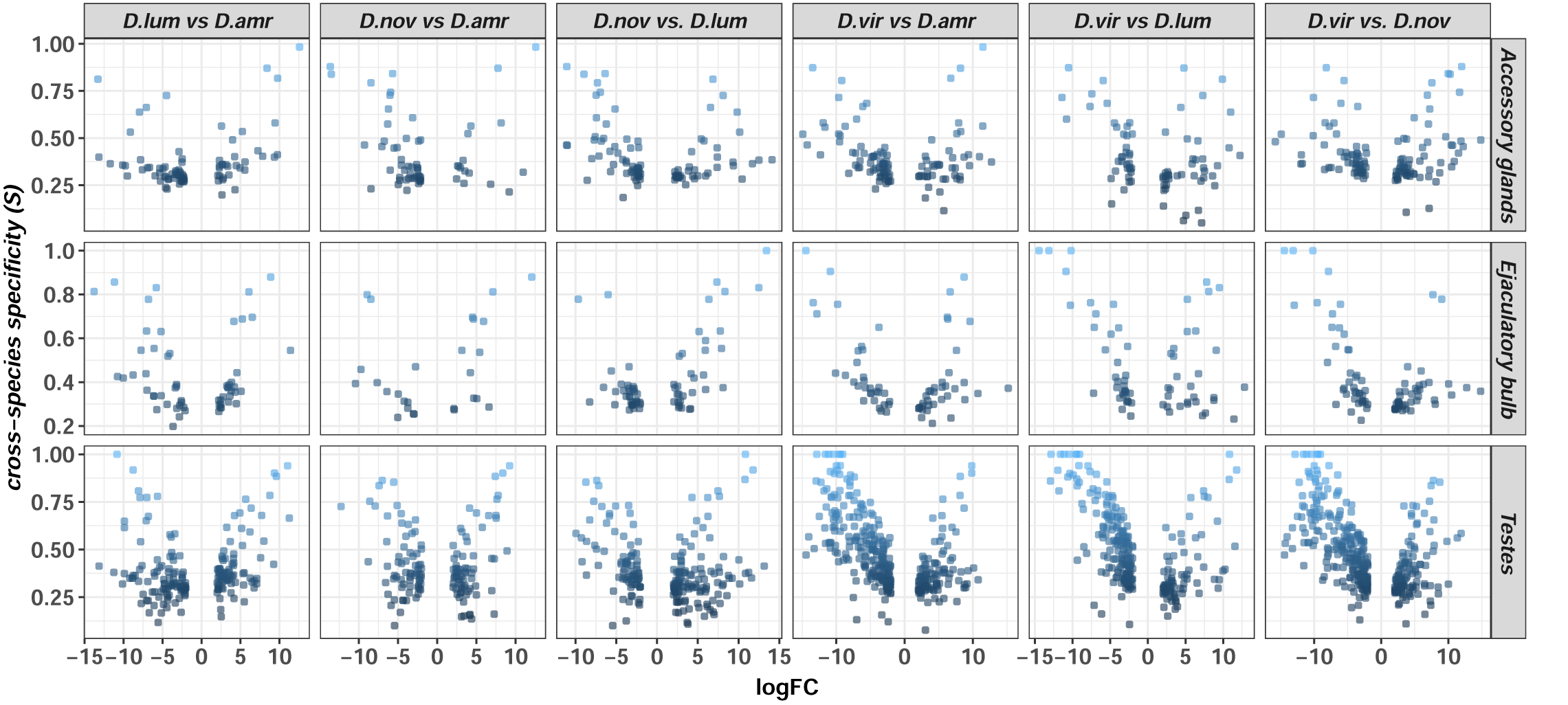
Tissue-biased differential abundance between species: The log-fold change (logFC) in pair-wise species comparisons of tissue-biased transcripts is plotted as a function of cross-species specificity (*S*). Genes with *S*>0.75 are highlighted.

**Figure S4.**
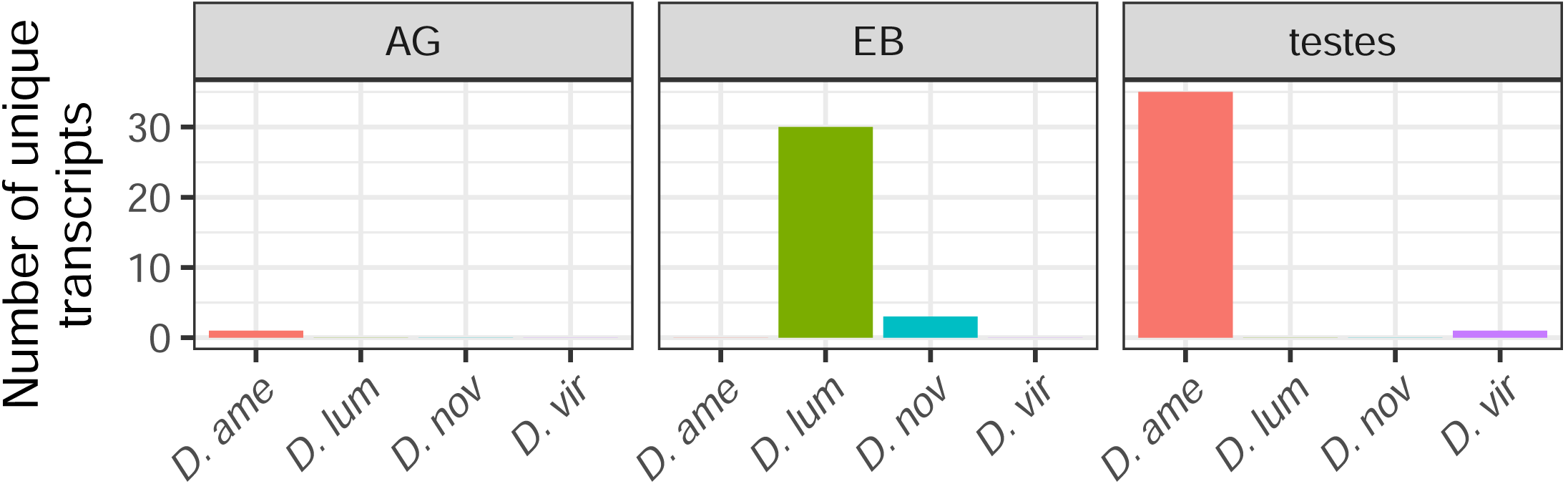
*De novo* transcripts: Number of transcripts in the *de novo* transcriptome assemblies of each species that have no detectable BLAST homology to transcripts in any of the sister species.

**Figure S5.**
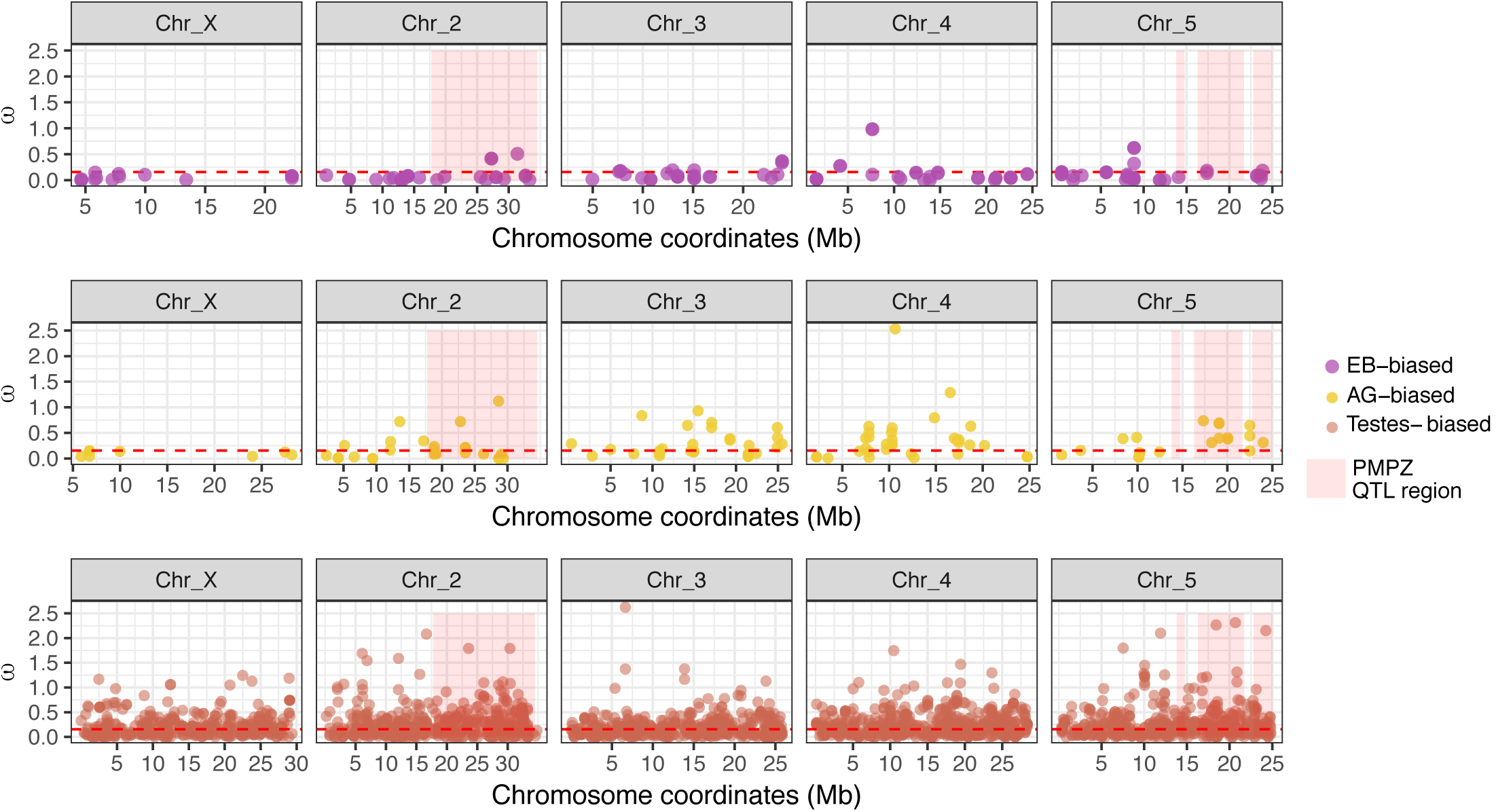
ω values of tissue-biased transcripts along the major chromosomes: ω values are plotted against genomic location for each tissue-biased transcripts category. The shaded pink regions highlight the paternal PMPZ QTLs identified previously (see text). The dashed red line indicates the genome average ω.

**Figure S6.**
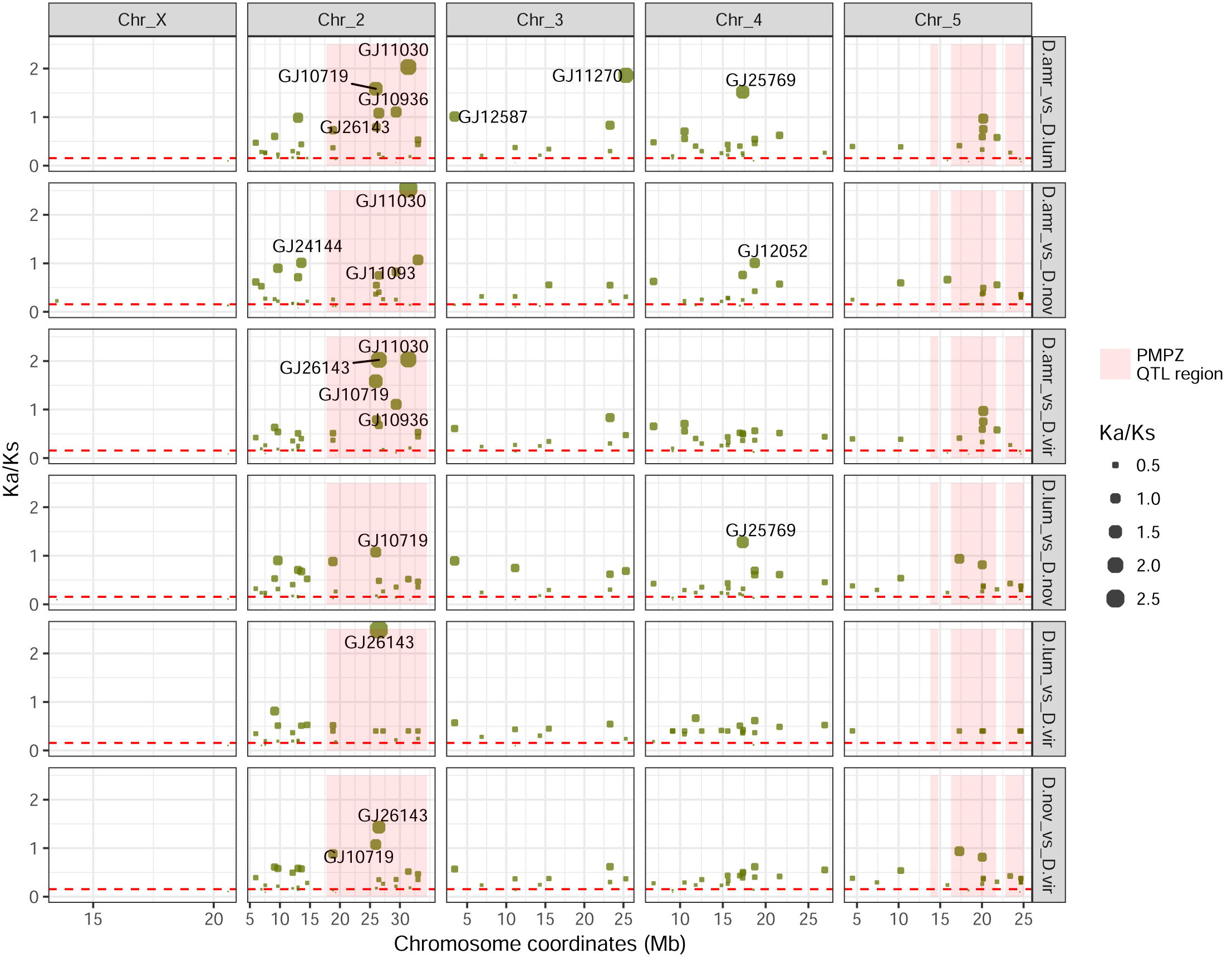
*K_a_/K_s_* of SFPs between species: Pairwise *K_a_/K_s_* values of AG-derived SFP candidates are plotted along the major chromosomes. Previously identified paternal PMPZ QTLs are shown in pink, and transcripts with *K_a_/K_s_* >1 are indicated. Point size corresponds to *y*-axis height (i.e. *K_a_/K_s_* value).

**Figure S7.**
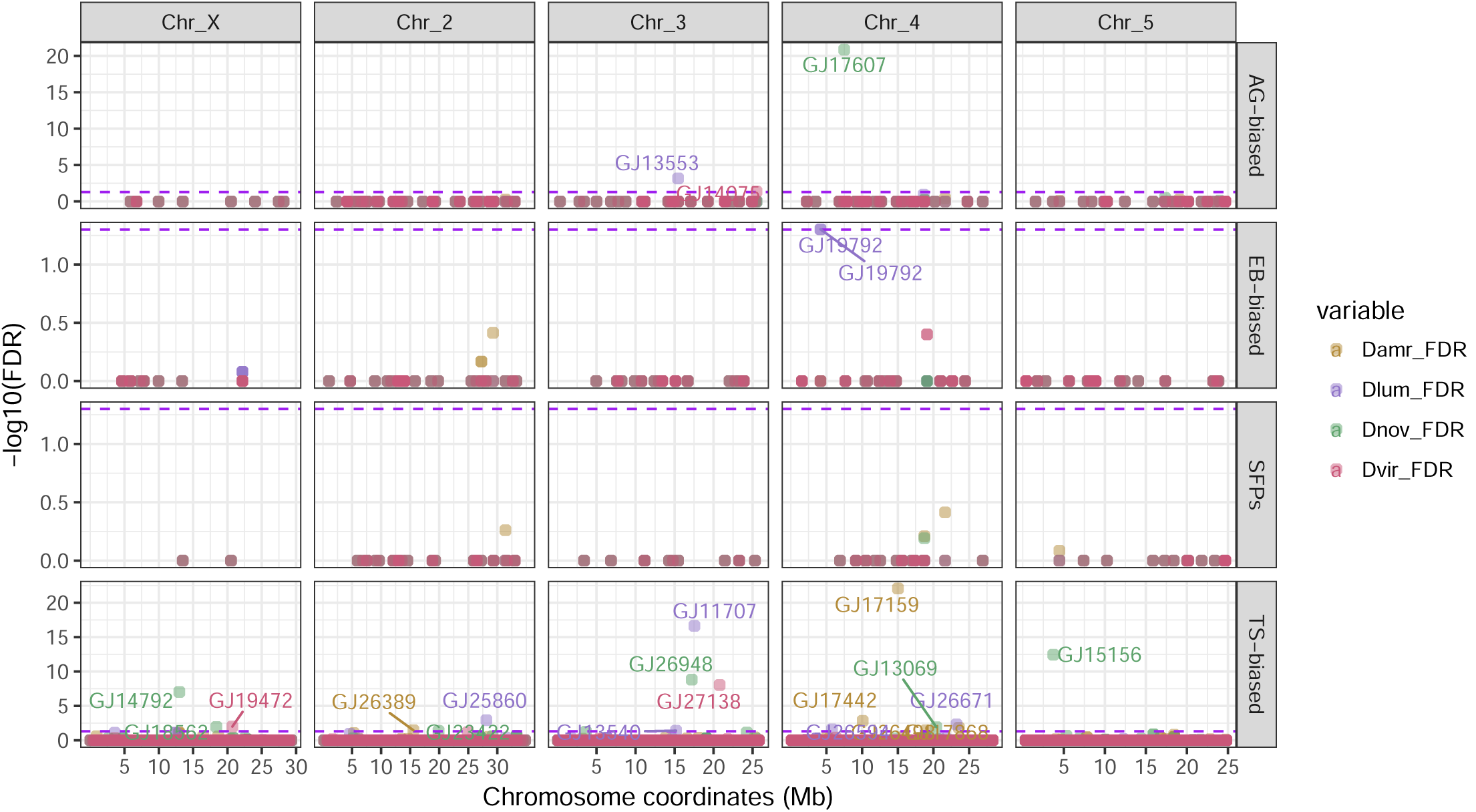
Distribution of transcripts with significant branch-site signature of positive selection: Genomic location of tissue-biased transcripts is plotted as a function of significance (–*log*_10_*FDR*) of the LRT for the branch-site test implemented in PAML. Point color corresponds to the terminal branch species, and the dotted line represents FDR = 0.05. Transcripts with *FDR<0.05* are indicated.

**Table S1.** GO enrichment among AG-biased and EB-biased genes (*FDR<*0.05)

| Gene class | Ontology | term | Number of genes | p-value |
| --- | --- | --- | --- | --- |
| AG-biased | Biological Process | protein glycosylation | 9 | 1.15E-06 |
|  |  | macromolecule glycosylation | 9 | 1.15E-06 |
|  |  | glycosylation | 9 | 1.42E-06 |
|  | Cellular Compartment | Golgi membrane | 10 | 1.25E-05 |
|  | Molecular Function | serine-type endopeptidase inhibitor activity | 9 | 3.04E-09 |
|  |  | endopeptidase inhibitor activity | 9 | 2.06E-08 |
|  |  | peptidase inhibitor activity | 9 | 2.65E-08 |
|  |  | endopeptidase regulator activity | 9 | 2.77E-08 |
|  |  | peptidase regulator activity | 9 | 7.41E-08 |
|  |  | enzyme inhibitor activity | 9 | 1.59E-06 |
|  |  | carbohydrate binding | 8 | 5.15E-06 |
| EB-biased | Biological Process | lipid metabolic process | 30 | 3.32E-17 |
|  |  | fatty acid biosynthetic process | 10 | 8.93E-11 |
|  |  | monocarboxylic acid metabolic process | 16 | 1.10E-10 |
|  |  | single-organism metabolic process | 43 | 2.18E-08 |
|  |  | oxoacid metabolic process | 18 | 2.11E-07 |
|  |  | carboxylic acid biosynthetic process | 10 | 2.54E-07 |
|  |  | organic acid metabolic process | 18 | 2.69E-07 |
|  |  | carboxylic acid metabolic process | 17 | 4.19E-07 |
|  |  | single-organism biosynthetic process | 21 | 5.77E-07 |
|  |  | prostanoid metabolic process | 4 | 7.88E-07 |
|  |  | small molecule biosynthetic process | 11 | 1.67E-06 |
|  |  | long-chain fatty-acyl-CoA metabolic process | 5 | 4.28E-06 |
|  |  | coenzyme metabolic process | 8 | 1.83E-05 |
|  |  | icosanoid metabolic process | 4 | 1.90E-05 |
|  |  | oxidation-reduction process | 16 | 1.96E-05 |
|  |  | fatty acid catabolic process | 5 | 1.97E-05 |
|  |  | wax biosynthetic process | 4 | 2.30E-05 |
|  |  | unsaturated fatty acid metabolic process | 4 | 2.84E-05 |
|  |  | fatty acid beta-oxidation using acyl-CoA oxidase | 3 | 2.88E-05 |
|  |  | steroid metabolic process | 7 | 3.94E-05 |
|  |  | cofactor metabolic process | 8 | 5.82E-05 |
|  |  | monocarboxylic acid catabolic process | 5 | 6.49E-05 |
|  |  | single-organism catabolic process | 13 | 8.95E-05 |
|  |  | lipid catabolic process | 7 | 0.000110127 |
|  | Cellular Component | microbody | 10 | 8.08E-08 |
|  |  | peroxisome | 9 | 2.57E-07 |
|  |  | integral component of endoplasmic reticulum membrane | 5 | 2.33E-05 |
|  | Molecular Function | cofactor binding | 12 | 4.51E-07 |
|  |  | coenzyme binding | 10 | 2.55E-06 |
|  |  | catalytic activity | 52 | 8.31E-06 |
|  |  | acyl-CoA dehydrogenase activity | 4 | 1.12E-05 |
|  |  | flavin adenine dinucleotide binding | 6 | 1.87E-05 |
|  |  | acyl-CoA oxidase activity | 3 | 1.99E-05 |
|  |  | long-chain-fatty-acyl-CoA reductase activity | 4 | 2.30E-05 |
|  |  | transferase activity, transferring acyl groups | 8 | 5.21E-05 |
|  |  | oxidoreductase activity | 17 | 5.39E-05 |
|  |  | sulfotransferase activity | 4 | 6.02E-05 |
|  |  | fatty-acyl-CoA binding | 4 | 8.35E-05 |
|  |  | transferase activity, transferring acyl groups other than amino-acyl groups | 7 | 0.00011877 |

**Table S2.** GO enrichment among testes-biased genes (*FDR*<0.001)

| Gene class | Ontology | Term | Number of genes | p-value |
| --- | --- | --- | --- | --- |
| Testes-biased | Biological Process | nuclear division | 78 | 2.43E-18 |
|  |  | microtubule-based movement | 50 | 1.29E-17 |
|  |  | organelle fission | 79 | 1.66E-17 |
|  |  | protein ubiquitination | 71 | 1.57E-13 |
|  |  | axonemal dynein complex assembly | 18 | 3.69E-13 |
|  |  | cell cycle process | 134 | 3.84E-12 |
|  |  | mitochondrial transport | 33 | 8.01E-12 |
|  |  | spermatogenesis | 41 | 7.48E-11 |
|  |  | single-organism organelle organization | 179 | 7.80E-11 |
|  |  | cellular protein metabolic process | 260 | 8.10E-11 |
|  |  | protein modification process | 204 | 1.21E-10 |
|  |  | organelle organization | 222 | 7.58E-10 |
|  |  | meiotic nuclear division | 27 | 4.91E-09 |
|  |  | macromolecule modification | 210 | 6.52E-09 |
|  |  | flagellated sperm motility | 16 | 1.22E-08 |
|  |  | male meiosis | 16 | 1.86E-08 |
|  |  | proteolysis involved in cellular protein catabolic process | 60 | 2.63E-08 |
|  |  | proteasomal protein catabolic process | 40 | 9.90E-08 |
|  |  | modification-dependent protein catabolic process | 51 | 1.51E-07 |
|  |  | protein catabolic process | 44 | 1.75E-07 |
|  |  | organelle assembly | 49 | 1.90E-07 |
|  |  | sperm individualization | 17 | 1.94E-07 |
|  |  | glycolytic process | 16 | 2.42E-07 |
|  |  | gamete generation | 62 | 2.92E-07 |
|  |  | citrate metabolic process | 19 | 4.02E-07 |
|  |  | nuclear envelope organization | 14 | 5.88E-07 |
|  |  | tricarboxylic acid metabolic process | 19 | 9.82E-07 |
|  |  | cell projection assembly | 28 | 1.85E-06 |
|  | Cellular Component | cilium | 69 | 2.40E-27 |
|  |  | cytoplasm | 362 | 7.37E-17 |
|  |  | microtubule associated complex | 84 | 8.94E-17 |
|  |  | centrosome | 43 | 6.68E-10 |
|  |  | Cul3-RING ubiquitin ligase complex | 18 | 1.43E-09 |
|  |  | intracellular part | 918 | 3.27E-09 |
|  |  | cytoplasmic dynein complex | 14 | 1.43E-07 |
|  |  | chromosome, centromeric region | 19 | 3.60E-07 |
|  |  | sperm part | 16 | 6.69E-07 |
|  |  | proteasome core complex | 16 | 7.79E-07 |
|  |  | mitochondrial membrane | 92 | 3.05E-06 |
|  |  | ribonucleoprotein granule | 25 | 8.16E-06 |
|  |  | sperm flagellum | 8 | 8.24E-06 |
|  | Molecular Function | cytoskeletal protein binding | 59 | 1.19E-08 |
|  |  | protein transmembrane transporter activity | 12 | 9.66E-08 |
|  |  | nucleotide binding | 220 | 1.52E-07 |
|  |  | ubiquitin-protein transferase activity | 41 | 4.03E-07 |
|  |  | purine nucleotide binding | 174 | 4.13E-07 |
|  |  | porin activity | 8 | 1.04E-06 |
|  |  | threonine-type endopeptidase activity | 16 | 1.32E-06 |
|  |  | nucleoside binding | 171 | 2.78E-06 |
|  |  | ribonucleoside binding | 169 | 3.91E-06 |
|  |  | purine ribonucleoside triphosphate binding | 167 | 6.80E-06 |

